# Identification and characterization of a SARS-CoV-2 M^pro^ G23 deletion ensitrelvir-resistant mutant

**DOI:** 10.1101/2025.09.30.679586

**Authors:** Yao Ma, Chengjin Ye, R. Rahisuddin, Sara H. Mahmoud, Anastasija Cupic, Ahmed Magdy Khalil, Esteban Castro, Nathaniel Jackson, Mahmoud Bayoumi, Yogesh K. Gupta, Adolfo Garcia-Sastre, Richard K Plemper, Luis Martinez-Sobrido

**Affiliations:** Texas Biomedical Research Institute, San Antonio, TX, USA; Greehey Children’s Cancer Research Institute, University of Texas Health Science Center, San Antonio, TX, USA; Department of Biochemistry and Structural Biology, University of Texas Health Science Center, San Antonio, TX, USA; Center of Scientific Excellence for Influenza Viruses, National Research Centre, Giza, Egypt; Graduate School of Biomedical Sciences, Icahn School of Medicine at Mount Sinai, New York, NY, USA; Department of Microbiology, Icahn School of Medicine at Mount Sinai, New York, NY, USA; Faculty of Veterinary Medicine, Zagazig University, Zagazig, Egypt; Virology Department, Faculty of Veterinary Medicine, Cairo University, Giza, Egypt; Global Health and Emerging Pathogens Institute, Icahn School of Medicine at Mount Sinai, New York, NY, USA; Department of Medicine, Icahn School of Medicine at Mount Sinai, New York, NY, USA; The Tisch Cancer Institute, Icahn School of Medicine at Mount Sinai, New York, NY, USA; Department of Pathology, Molecular and Cell-Based Medicine, Icahn School of Medicine at Mount Sinai, New York, NY, USA; The Icahn Genomics Institute, Icahn School of Medicine at Mount Sinai, New York, NY, USA; Center for Translational Antiviral Research, Institute for Biomedical Sciences, Georgia State University, Atlanta, Georgia, USA

**Keywords:** SARS-CoV-2, Ensitrelvir, attenuated virus, drug-resistance mutations, G23 deletion

## Abstract

Ensitrelvir is an antiviral drug that specifically targets the main protease (M^pro^) of Severe Acute Respiratory Syndrome Coronavirus 2 (SARS-CoV-2) has been approved for the treatment of coronavirus disease 2019 (COVID-19) due to the conservation of its target protein which is essential in the viral lifecycle. However, SARS-CoV-2 could introduce mutations in the viral proteins to confer resistance to antivirals. Thus, screening for drug-resistant SARS-CoV-2 mutants and elucidating their resistant mechanisms are critical for guiding the selection of effective antiviral therapies. Here, we utilized a luminescent attenuated SARS-CoV-2 (Δ3a7b-Nluc WT) to safely identify ensitrelvir drug-resistant mutants (DRM-E) without the need of using virulent forms of SARS-CoV-2. We isolated a DRM-E containing a G23 deletion (G23del) in M^pro^ with high resistance (>1,000 fold) to ensitrelvir, but not to the other M^pro^ inhibitor (nirmatrelvir) or to the RNA-dependent RNA polymerase (RdRp) inhibitor remdesivir. The contribution of G23del was confirmed by generating a recombinant luminescent attenuated SARS-CoV-2 containing G23del in the non-structural protein 5 (NSP5) gene (Δ3a7b-Nluc G23del). Δ3a7b-Nluc G23del exhibited significant resistance to ensitrelvir in both cultured cells an in K18 hACE2 transgenic mice. Binding affinity revealed that G23del mutation substantially altered M^pro^ binding affinity for ensitrelvir but not nirmatrelvir. In conclusion, our results demonstrate that G23del in M^pro^ can confer high resistance to ensitrelvir. Positively, G23del in M^pro^ does not render SARS-CoV-2 resistant to nirmatrelvir or remdesivir, suggesting the feasibility of treating infections with SARS-CoV-2 containing G23del with these other approved antivirals.

**SIGNIFICANCE:** The clinical use of SARS-CoV-2 antiviral drugs is increasingly challenged by the emergence of drug-resistant mutants. Thus, there is a pressing need to identify and characterize antiviral escape SARS-CoV-2 variants, particularly for FDA-approved antivirals. Our study addresses this by employing a luminescent attenuated virus platform (Δ3a7b-Nluc WT) to safely identify and characterize resistance mutations without the concern of using virulent forms of SARS-CoV-2. Using this safe approach, we have identified a G23 deletion (G23del) in SARS-CoV-2 M^pro^, which mediates resistance to ensitrelvir *in vitro* and *in vivo*. Importantly, while G23del was able to confer more than 1,000-fold increased resistance to ensitrelvir, SARS-CoV-2 containing G23del remained sensitive to other M^pro^ (nirmatrelvir) and RdRp (remdesivir) inhibitors. Altogether, this study demonstrates the feasibility of using Δ3a7b-Nluc to safely identify and characterize drug resistant viruses without the biosafety concern of using virulent SARS-CoV-2 and advance the design of next-generation antiviral drugs.

## INTRODUCTION

Over the past five years, the coronavirus disease 2019 (COVID-19) pandemic has resulted in approximately seven million deaths worldwide (1). In addition to its significant global health impact, the COVID-19 pandemic has imposed substantial economic burdens, further compounded by the continuous evolution of severe acute respiratory syndrome coronavirus 2 (SARS-CoV-2) (2–5). This continuous viral evolution has diminished the efficacy of vaccines developed against the original strain, necessitating continual vaccine reformulation to maintain protective immunity against emerging variants (6–10). Similarly, monoclonal antibodies initially granted emergency use authorization (EUA) in the United States (US) have rapidly lost neutralization potency against newly emerging SARS-CoV-2 variants, culminating in the revocation of all clinical use authorizations by early 2023 (11–13). In contrast, small-molecule antiviral drugs have largely retained efficacy owing to their targeting of highly conserved proteins essential to the viral life cycle. However, the emergence of drug-resistant mutants poses a growing challenge to the clinical effectiveness against SARS-CoV-2 (14–19). Therefore, it is crucial to identify and characterize SARS-CoV-2 resistant variants to currently approved antivirals. These efforts, which elucidate escape mechanisms and impacts on drug efficacy, are necessary to guide therapeutic strategies and advance the development of next-generation antivirals for the treatment of SARS-CoV-2 and potentially newly emerging coronaviruses.

As a cysteine protease encoded by the viral nonstructural protein 5 (NSP5) gene, the main protease (M^pro^) plays a critical role in the SARS-CoV-2 life cycle by cleaving the viral polyproteins derived from open reading frame (ORF) 1a and 1b proteins at 11 conserved sites to generate 12 functional NSP essential for viral genome replication and gene transcription (20, 21). Owing to its indispensable function and high degree of sequence conservation among β-coronaviruses, M^pro^ is a promising target for broad-spectrum antivirals (22). The non-covalent, non-peptide inhibitor ensitrelvir, which was first approved in Japan, binds to the dimeric form of M^pro^ by specifically recognizing its S1, S2, and S1′ subsites, thereby inhibiting protease processing (23–25). Ensitrelvir antiviral activity has been demonstrated in cell culture and animal models, including prevention of direct viral transmission (26–28). Clinical trials have proved that ensitrelvir exhibits favorable pharmacokinetic properties, including a prolonged half-life, and is well-tolerated in humans (29). Furthermore, it has been shown that ensitrelvir is highly effective at reducing viral load in patients with mild-to-moderate COVID-19, leading to its emergency use authorization by the US Food and Drug Administration (FDA) (30–32). However, mutations conferring resistance to ensitrelvir have been identified within NSP5 gene, including D48G, M49I/L, P52S, S144A, E166A/V, L167F, P168del, as well as the combination M48L+S144 and M49L+E166A, all of which reduce the antiviral potency of ensitrelvir *in vitro* (33–35). Critically, an *in vivo* study demonstrated that ensitrelvir treatment was ineffective in hamsters infected with a virus harboring the M49L+166A mutations (18). Consequently, close surveillance of the emergence of ensitrelvir-resistant variants is essential to inform and guide appropriate antiviral treatment strategies.

An attenuated, luminescent recombinant SARS-CoV-2 strain (Δ3a7b-Nluc WT) has been engineered through the deletion of the accessory ORFs 3a and 7b and the incorporation of a nanoluciferase (Nluc) reporter gene (36). This recombinant Δ3a7b-Nluc WT was designed to establish a safer platform for the identification and characterization of drug-resistant mutants, circumventing the biosafety concerns associated with using virulent forms of SARS-CoV-2. While ORFs 3a and 7b deletions contribute to viral attenuation, Nluc expression enables sensitive, real-time tracking of viral replication (37–39), providing a robust and safe approach to select and validate mutations responsible for SARS-CoV-2 drug resistance to antivirals.

In this study, we used Δ3a7b-Nluc WT to identify ensitrelvir drug-resistant mutants (DRM-E). By serially passaging Δ3a7b-Nluc WT in Vero cells in the presence of increasing concentrations of ensitrelvir, we identified a DRM-E exhibiting high resistance (>1,000 fold) to ensitrelvir without affected its sensitivity to other M^pro^ inhibitor (nirmatrelvir) or to the RNA-dependent RNA polymerase (RdRp) inhibitor remdesivir. Next-generation sequencing of DRM-E revealed a glycine deletion at position 23 (G23del) in M^pro^, which is encoded by the nonstructural protein 5 (NSP5) gene. To demonstrate the contribution of G23del to DRM-E resistance to ensitrelvir, we engineered a recombinant Δ3a7b-Nluc harboring the G23del mutation (Δ3a7b-Nluc G23del). Δ3a7b-Nluc G23del recapitulated the high resistance to ensitrelvir both *in vitro* and *in vivo* observed with DRM-E. Importantly, a quantitative measurement of the binding affinities indicated that G23del substantially reduced ensitrelvir’s binding to the M^pro^ active site without affecting binding of nirmatrelvir. Collectively, these findings demonstrate the importance of viral surveillance for emerging protease mutations to inform clinical antiviral decision-making. Our findings also demonstrate the feasibility of using Δ3a7b-Nluc WT to safely identify and characterized in laboratory settings SARS- CoV-2 drug resistant strains without the biosafety concern of conducting these experiments with wild-type forms of SARS-CoV-2. Finally, our findings demonstrate that G23del in M^pro^ significantly affect the antiviral activity of ensitrelvir to SARS-CoV-2.

## RESULTS

### Isolation of ensitrelvir drug resistant mutant (DRM-E)

To isolate viruses resistant to ensitrelvir, Δ3a7b-Nluc WT was serially passaged in Vero-AT cells in the presence of increasing concentrations of ensitrelvir (Figure 1A). Following 10 serial passages, a variant with significantly enhanced resistance, designated DRM-E, was selected. We assessed the resistance profiles of the parental P0 Δ3a7b-Nluc WT and the P10 DRM-E viruses by immunofluorescence assay (IFA) across a range of ensitrelvir concentrations (Figure 1B). At 0.03 µM ensitrelvir, P0 replication was only partially inhibited compared to the no-drug control (0 µM), whereas DRM-E replication was unaffected. At 3 µM ensitrelvir, we observed almost complete inhibition of P0 Δ3a7b-Nluc WT, whereas DRM-E maintained exhibited significant replication. Both viruses, P0 Δ3a7b-Nluc WT and P10 DRM-E, were fully inhibited using 300 µM ensitrelvir (Figure 1B). This result confirmed that P10 DRM-E had acquired substantial resistance to ensitrelvir.

**Figure 1.**
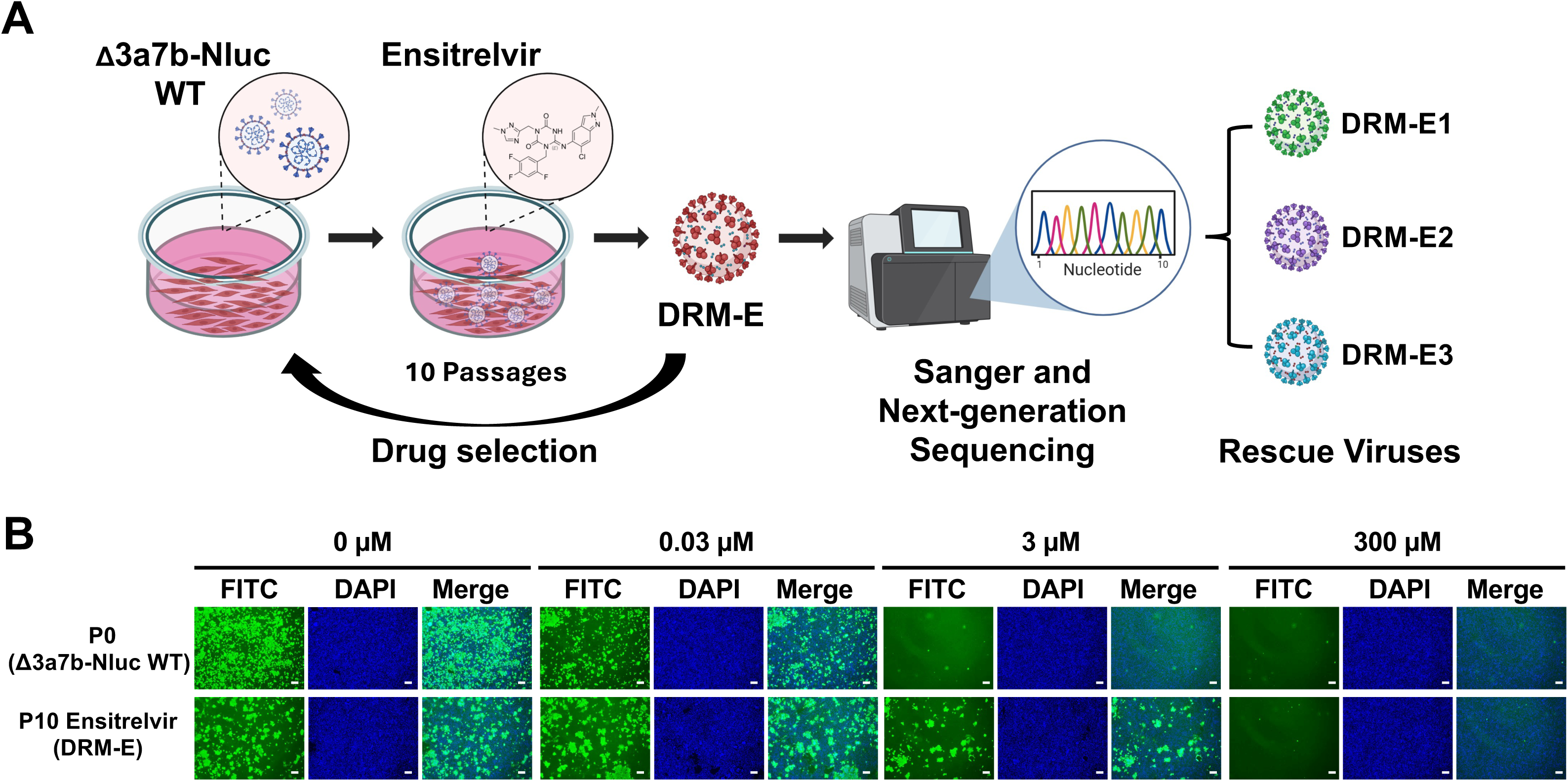
Selection and characterization of ensitrelvir-resistant mutant (DRM-E). A) Schematic representation of selection of DRM-E: The recombinant Δ3a7b-Nluc WT was serially passaged in Vero-AT cells under increasing concentrations of ensitrelvir for a total of 10 passages. RNA isolated from the P10 population (DRM-E) was subjected to next-generation sequencing (NGS), and Sanger sequencing to identify resistance-associated mutations. Mutations identified were introduced into the Δ3a7b-Nluc WT backbone using BAC-based reverse genetics for subsequent characterization. B) Δ3a7b-Nluc WT and DRM-E resistance to ensitrelvir: Vero-AT cells were infected (MOI = 0.001) with either P0 Δ3a7b-Nluc WT (top) or P10 ensitrelvir (DRM-E) (bottom). After viral adsorption, the inoculum was replaced with post-infection medium containing the indicated concentrations of nirmatrelvir. At 48 hours post-infection, cells were fixed, permeabilized, and subjected to immunofluorescence staining using a monoclonal nucleocapsid (N) protein antibody (1C7C7) and an FITC-conjugated secondary anti-mouse antibody. Nuclei were counterstained with DAPI. Scale bars = 200 µm.

To quantify the resistance of DRM-E to ensitrelvir and to assess its ability to be inhibited by a different M^pro^ inhibitor (nirmatrelvir) or a RdRp inhibitor (remdesivir), we determined the half-maximal effective concentration (EC_50_) using a plaque reduction neutralization test (PRNT) assay (Figure 2). DRM-E exhibited an ∼2,000-fold increase in EC_50_ against ensitrelvir compared to the parental Δ3a7b-Nluc WT virus (DRM-E EC_50_ = 56.49 µM compared to Δ3a7b-Nluc WT EC_50_ = 24.95 nM) (Figures 2A and 2D). Resistance was specific to ensitrelvir, as DRM-E showed no increased resistance to nirmatrelvir (DRM-E EC_50_ = 1.78 µM compared to Δ3a7b-Nluc WT EC_50_ = 1.38 µM) (Figures 2B and 2D) and remdesivir (DRM-E EC_50_ = 3.12 µM compared to Δ3a7b-Nluc WT EC_50_ = 4.90 µM) (Figures 2C and 2D) demonstrating that the acquired resistance of DRM-E is specific to ensitrelvir.

**Figure 2.**
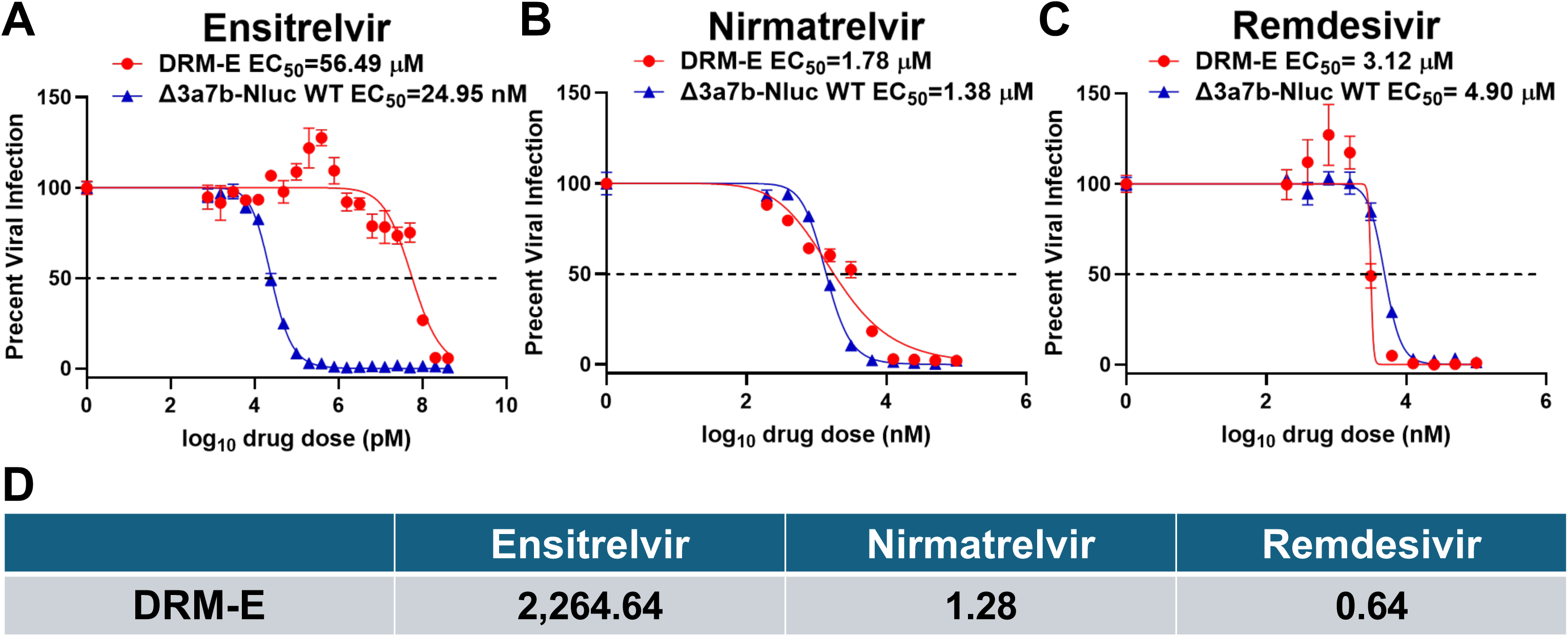
Drug susceptibility profile of the DRM-E variant. The antiviral activity of ensitrelvir (A), nirmatrelvir (B), and remdesivir (C) against parental P0 (blue) and DRM-E (red) were evaluated in Vero-AT cells using a plaque reduction neutralization test (PRNT). The half-maximal effective concentration (EC_₅₀_) values were calculated using nonlinear regression in GraphPad Prism. Data are presented as mean ± SD. The dotted line indicates 50% viral inhibition. Fold-increase in EC_₅₀_ values of DRM-E relative to the parental P0 virus for each antiviral agent is shown (D).

### Identification of DRM-E mutation responsible for resistance to ensiltrelvir

To identify the mutation(s) conferring ensitrelvir resistance in the DRM-E mutant, we performed next-generation sequencing (NGS) on RNA sample collected from Vero-AT cells infected with DRM-E mutant (Figure 3A). As an internal control, we also sequenced a virus that was serially passaged 10 times in the absence of ensitrelvir (P10 PBS). NGS analysis revealed four amino acid mutations location in the viral NSP present at a frequency >30% in the control P10 PBS population: NSP3 (E1270D, 99.8%), NSP4 (A260V, 99.7%, and M458I, 50.0%), and NSP15 (G13E, 99.9%). In contrast, the DRM-E population harbored six mutations in NSP: NSP5 (G23 deletion, 84.2%), NSP12 (N447 deletion, 62.0%), NSP13 (N95T, 82.2%), NSP14 (P46S, 76.6%, and G59S, 69.9%), and NSP15 (S97A, 76.6%). Since ensitrelvir specifically targets the main protease encoded by NSP5, the G23 deletion (G23del) within NSP5 gene was prioritized as the most likely candidate responsible for the resistant phenotype. To validate the presence of G23del in the NSP5 gene of DRM-E, we performed RT-PCR and Sanger sequencing, which confirmed the presence of the G23del (Figure 3B). Structural mapping localized this mutation to a β-hairpin loop (residues T21–T26) that forms part of the S1’ subsite of the M^pro^ substrate-binding pocket (Figure 3C), a critical region for ensitrelvir binding.

**Figure 3.**
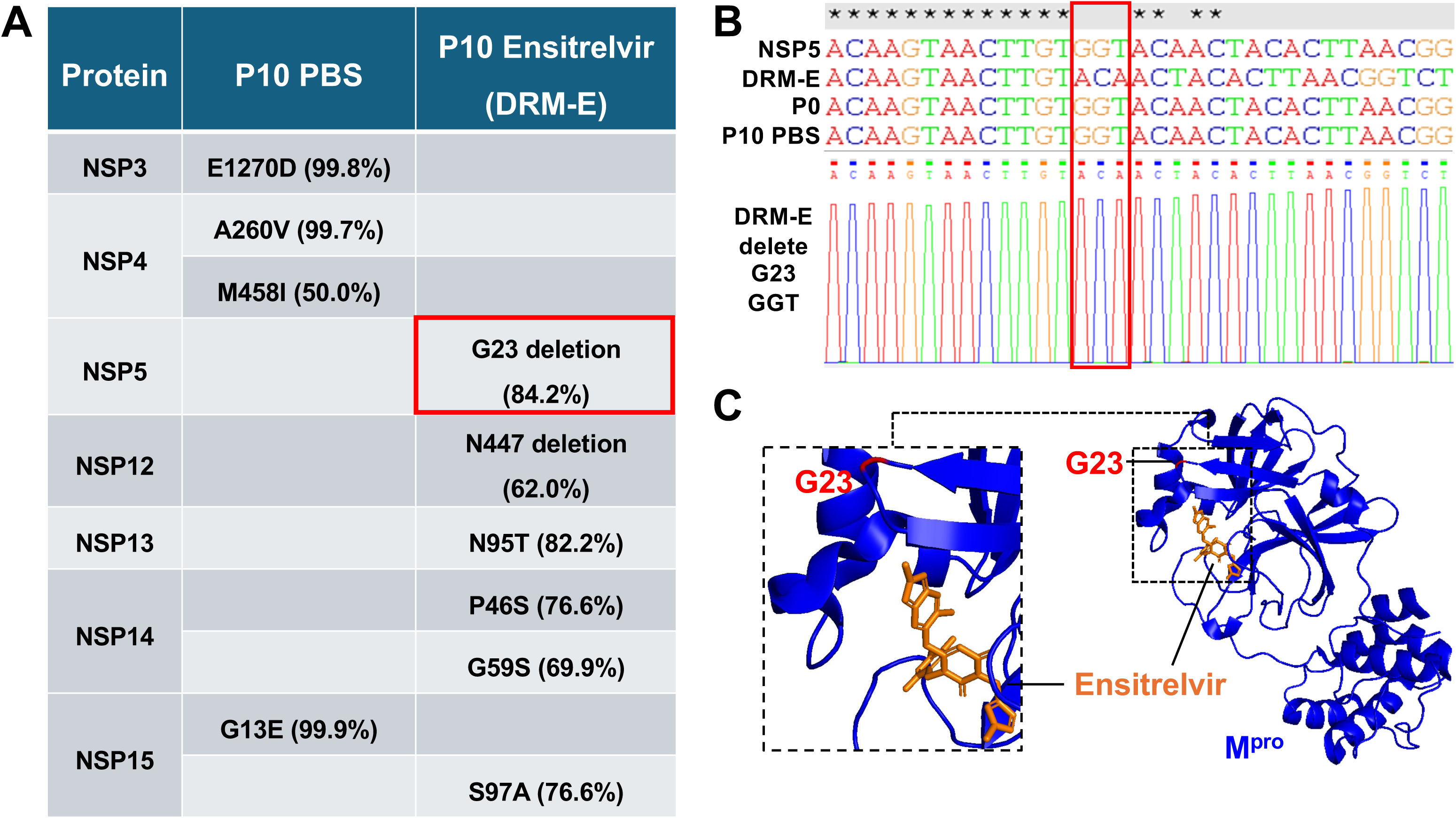
Identification of resistance-conferring mutations in the DRM-E variant. A) NGS was performed on RNA isolated from Δ3a7b-Nluc WT populations serially passaged: in the presence of PBS (P10 PBS) or increasing concentrations of ensitrelvir (P10 Ensitrelvir, DRM-E). Amino acid differences at a frequency >30% are listed with their respective location in the viral NSP. The G23 deletion (G23del) in NSP5, a candidate mutation responsible for resistance to ensitrelvir, is highlighted with a red rectangle. B) The NSP5 coding region of the same RNA samples were amplified by RT-PCR and analyzed by Sanger sequencing. The G23del mutation in DRM-E is indicated. C) Amino acid G23 (red) in M^pro^ (blue) is shown. Ensitrelvir is indicated in orange. The M^pro^-ensitrelvir complex was obtained from PDBID: 8HBK.

### Characterization of M^pro^ G23del mutation identified in DRM-E

To investigate the contribution of the M^pro^ G23del to ensitrelvir resistance, we engineered a recombinant Δ3a7b-Nluc G23del virus harboring this mutation. Plaque assays in Vero-AT cells revealed that the Δ3a7b-Nluc G23del mutant formed smaller plaques than the parental Δ3a7b-Nluc WT (Figure 4A), suggesting a potential impact on viral spread and/or relative fitness in cell culture. Subsequent growth analysis demonstrated that the Δ3a7b-Nluc G23del mutant was slightly attenuated compared to the Δ3a7b-Nluc WT virus, with lower titers at 24- and 48-hours post-infection (Figure 4B). This replication defect was corroborated by slightly delayed Nluc expression kinetics (Figure 4C), confirming that G23del mutation carries a fitness penalty *in vitro*. Notably, viral titers and levels of Nluc at 72 hours post-infection of Δ3a7b-Nluc G23del were comparable to those of Δ3a7b-Nluc WT (Figures 4B and 4C, respectively).

**Figure 4.**
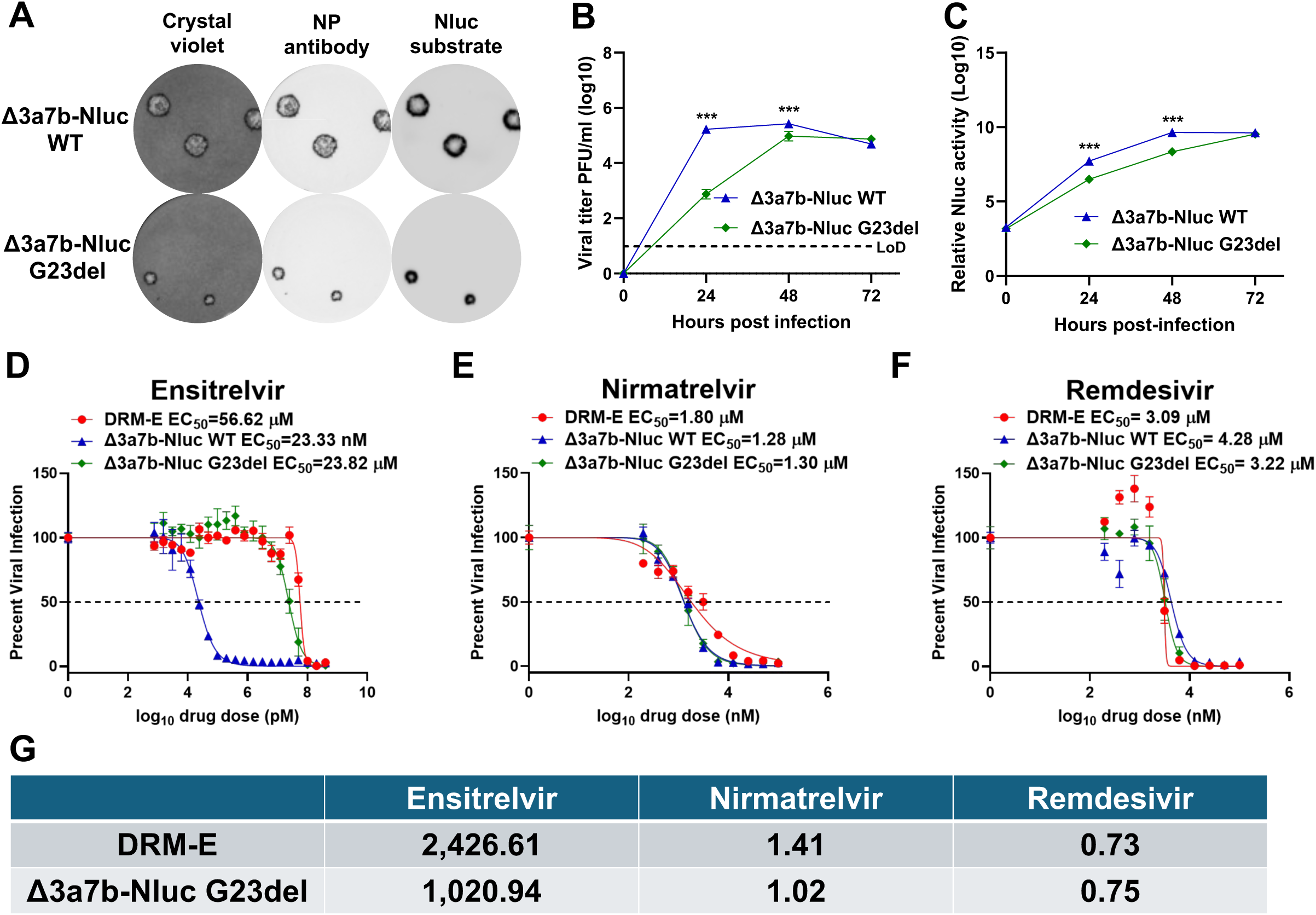
Characterization of Δ3a7b-Nluc G23del mutant. A) Plaque assay: Viral plaque morphology of Δ3a7b-Nluc WT (top) and Δ3a7b-Nluc G23del mutant (bottom) in Vero-AT cells. Viral plaques were visualized by crystal violet staining (left), immunostaining with the anti-N protein antibody 1C7C7 (middle), or Nluc substrate (right). B-C) Growth kinetics and nluc expression of Δ3a7b-Nluc WT and Δ3a7b-Nluc G23del viruses: Vero-AT cells were infected at an 0,01 MOI. Cell culture supernatants were collected at the indicated time points, and viral titers were quantified by plaque assay (B), and Nluc activity (C). D-F) Drug susceptibility profiles: The antiviral activity of ensitrelvir (D), nirmatrelvir (E), and remdesivir (F) against Δ3a7b-Nluc WT, DRM-E, and Δ3a7b-Nluc G23del was determined by PRNT assay. EC_₅₀_ values were calculated using GraphPad Prism. (G) Fold-change in EC_₅₀_ values for DRM-E and Δ3a7b-Nluc G23del relative to Δ3a7b-Nluc WT. Data represent mean ± SD. The dotted line indicates 50% viral inhibition. Statistical significance was determined by one-way ANOVA with Tukey’s post hoc test (*p < 0.05, **p < 0.01, ***p < 0.001).

We next evaluated the susceptibility profile of the recombinant Δ3a7b-Nluc G23del mutant to ensitrelvir. Δ3a7b-Nluc G23del exhibited a pronounced (∼1,000-fold) increase in the EC_₅₀_ for ensitrelvir (EC_₅₀_ = 23.82 µM) compared to the parental Δ3a7b-Nluc WT (EC_₅₀_ = 23.33 nM) (Figures 4D and 4G). In contrast, the EC_₅₀_ values of Δ3a7b-Nluc G23del for nirmatrelvir (EC_50_ = 1.30 µM) and remdesivir (EC_50_ = 3.22 µM) remained unchanged and were comparable to those of Δ3a7b-Nluc WT and the DRM-E (Figures 4E-4G). These results demonstrate that the G23del mutation in M^pro^ is responsible for the high resistance of DRM-E to ensitrelvir.

Next, we assessed the *in vivo* efficacy of ensitrelvir against the Δ3a7b-Nluc G23del variant using the K18-hACE2 mouse model of SARS-CoV-2 infection (Figure 5A). Mice were infected intranasally with 10^7^ PFU of either Δ3a7b-Nluc WT or Δ3a7b-Nluc G23del. After one day post-infection, mice were treated with ensitrelvir (60 mg/kg, twice daily) for three continuous days. Nluc-derived luminescence in lung and viral load in the lung and nasal turbinate of infected mice were quantified on day 4 post-infection. In mice infected with the Δ3a7b-Nluc WT, ensitrelvir treatment resulted in a significant reduction of Nluc expression in the lungs (Figure 5B) and a corresponding significant decrease in viral titers in both the lungs and nasal turbinate (Figures 5C and 5D, respectively). In contrast, ensitrelvir treatment of mice infected with the Δ3a7b-Nluc G23del yielded no therapeutic benefit. No significant reduction in Nluc expression was observed in the lungs of mice infected with Δ3a7b-Nluc G23del in the presence of ensitrelvir (Figure 5B), and viral titers in the lungs and nasal turbinate of mice treated with ensitrelvir were statistically indistinguishable from those in the untreated control group (Figures 5E and 5F). These data demonstrate that M^pro^ G23del confers resistance to ensitrelvir *in vivo*, abolishing its antiviral efficacy.

**Figure 5.**
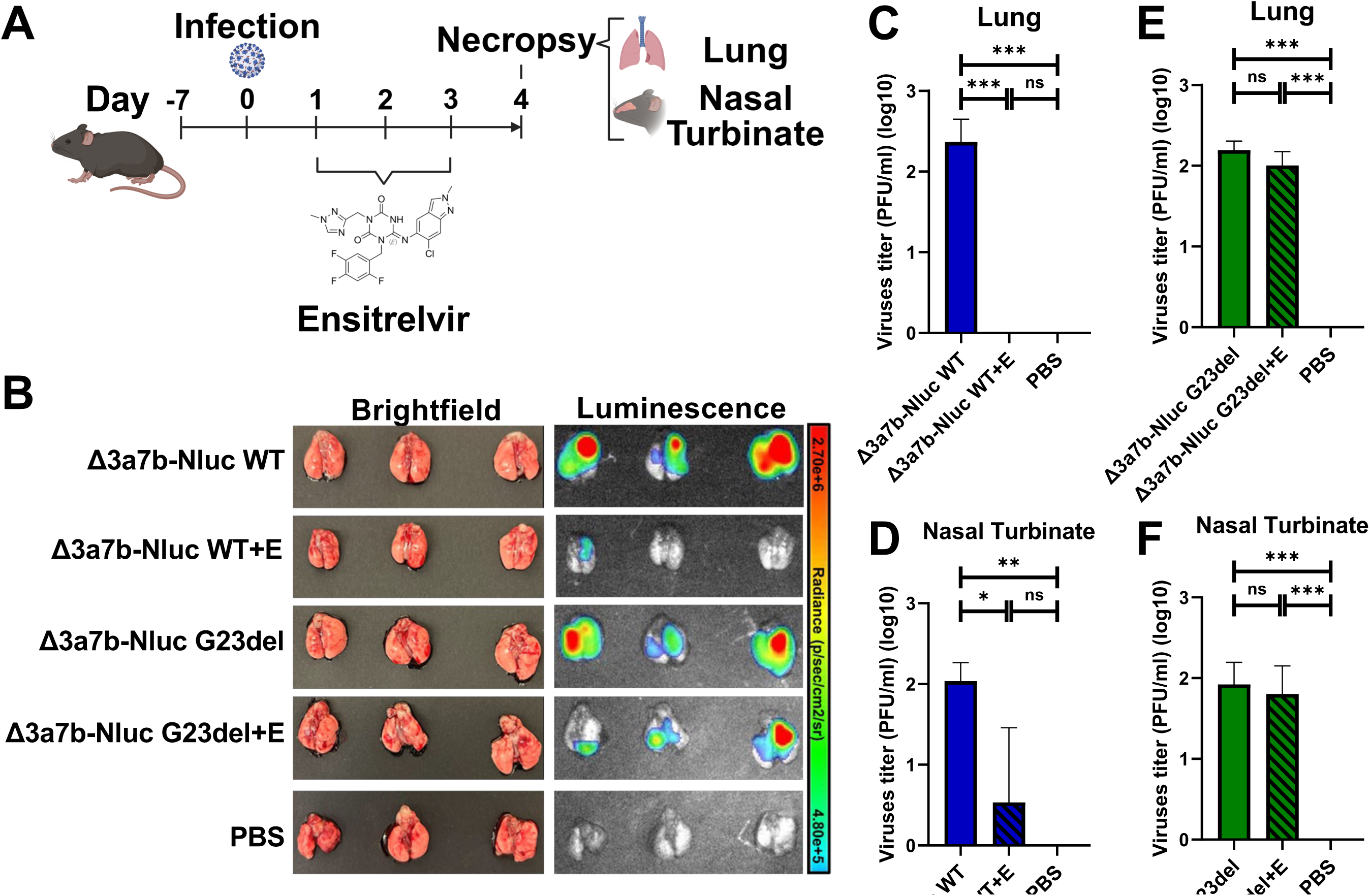
*In vivo* efficacy of ensitrelvir. A) Schematic representation of the experimental timeline of ensitrelvir antiviral therapy in K18-hACE2 transgenic mice infected with either Δ3a7b-Nluc WT or the Δ3a7b-Nluc G23del mutant. B) Nluc-derived luminescence in lungs using an *in vivo* imaging system (IVIS). C-F) Viral titrations: following infection or antiviral treatment. Viral loads in homogenates of lungs (C, E) and nasal turbinates (D, F) from K18-hACE2 mice at 4 days post-infection, with or without ensitrelvir treatment, were determined by plaque assay.

### Detection of ensitrelvir-specific resistance

The binding affinities of ensitrelvir and nirmatrelvir to M^pro^-WT and M^pro^-G23del purified proteins were determined using isothermal titration calorimetry (ITC) under identical experimental conditions. The ITC experiments were performed at two different protein-ligand concentrations: low (20 µM protein and 0.2 mM ligand) (Figure 6) and high (30 µM ligand and 0.3 mM) (Figure S1). At a low concentration, the equilibrium dissociation constant (K_d_) for ensitrelvir binding to M^pro^-WT was 333 ± 63 nM (Figure 6A). Interestingly, the deletion of G23 completely abolished the binding of ensitrelvir to M^pro^-G23del protein under identical experimental settings (Figure 6B). At the same protein and ligand concentration, the K_d_ for nirmatrelvir binding to M^pro^-WT was 144 ± 47 nM (Figure 6C), whereas for M^pro^-G23del was ∼2.5-fold lower (K_d_ = 377 ± 67 nM; Figure 6D). Similar results were observed at high protein and ligand concentrations (Figure S1). Consistently. ensitrelvir showed a K_d_ of 297 ± 203 nM for M^pro^-WT and no binding with M^pro^-G23del mutant (Figures S1A and S1B). However, Nirmatrelvir showed slightly better binding to M^pro^-G23del (K_d_ = 209 ± 68 nM) compared to M^pro^-WT (K_d_ = 274 ± 99 nM) at this concentration (Figures S1C and S1D). These results corroborate our *in vitro* and *in vivo* findings demonstrating a lack of activity of ensitrelvir but not nirmatrelvir against M^pro^-G23del. We noted a discrepancy in K_d_ values for ensitrelvir and nirmatrelvir to the M^pro^-WT in our results (K_d_ = 144 – 333 nM) and that previously reported in the literature (K_d_ = 6 - 8 nM), which is mainly due to the presence of two additional amino acids (GS) in our M^pro^ (25, 40, 41). Since we carried out the ITC experiments for both M^pro^-WT and G23del mutant under identical experimental conditions, including the presence of these two additional amino acid mutations, the relative difference in binding affinity supports our conclusion that G23 deletion abolishes the binding of ensitrelvir, but not nirmatrelvir to M^pro^.

**Figure 6.**
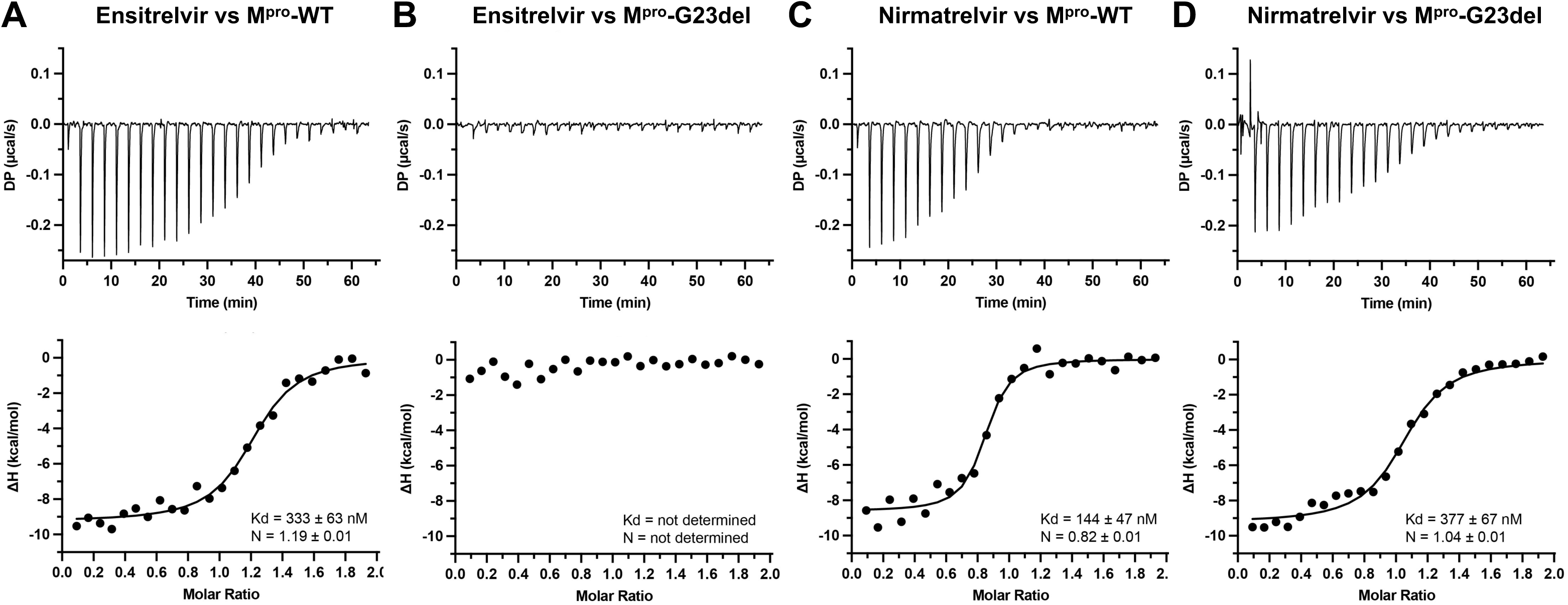
Binding isotherms of Ensitrelvir and Nirmatrelvir. Isothermal titration calorimetry (ITC) experiments with M^pro^-WT and M^pro^-G23del were performed using 20 µM protein and 0.2 mM of ligand at 25°C in identical experimental conditions. The upper panel illustrates the raw heat signals, while the lower panel shows the integrated heat and the fit using a one-site binding model for Ensitrelvir with M^pro^-WT (A), Ensitrelvir with M^pro^-G23del (B), Nirmatrelvir wtih M^pro^-WT (C), and Nirmatrelvir with M^pro^-G23del (D).

To further elucidate the mechanism by which deletion at G23 confers resistance to ensitrelvir, we employed AlphaFold2 to predict the three-dimensional structures of both M^pro^-WT and M^pro^-G23del (Figure 7). Structural models were aligned and analyzed using PyMOL. This analysis revealed that the primary structural alteration is predicted to occur within the β-hairpin loop (residues T21-T26) that forms part of the S1’ subsite of the substrate-binding pocket. The deletion of G23 was predicted to disrupt the conformation of this β-sheet. Specifically, ensitrelvir binding in this pose is dependent on its 6-chloro-2-methyl-2H-indazole moiety forming a critical hydrogen bond with the T26 in M^pro^-WT. The G23del mutation was posited to induce a structural shift that likely perturbed the orientation of T26, thereby compromising this essential interaction and reducing binding affinity. In contrast, the binding mode of nirmatrelvir does not involve interactions with this β-hairpin loop, providing a structural basis for the drug resistance specificity of the G23del mutation, conferring resistance to ensitrelvir but not to nirmatrelvir.

**Figure 7.**
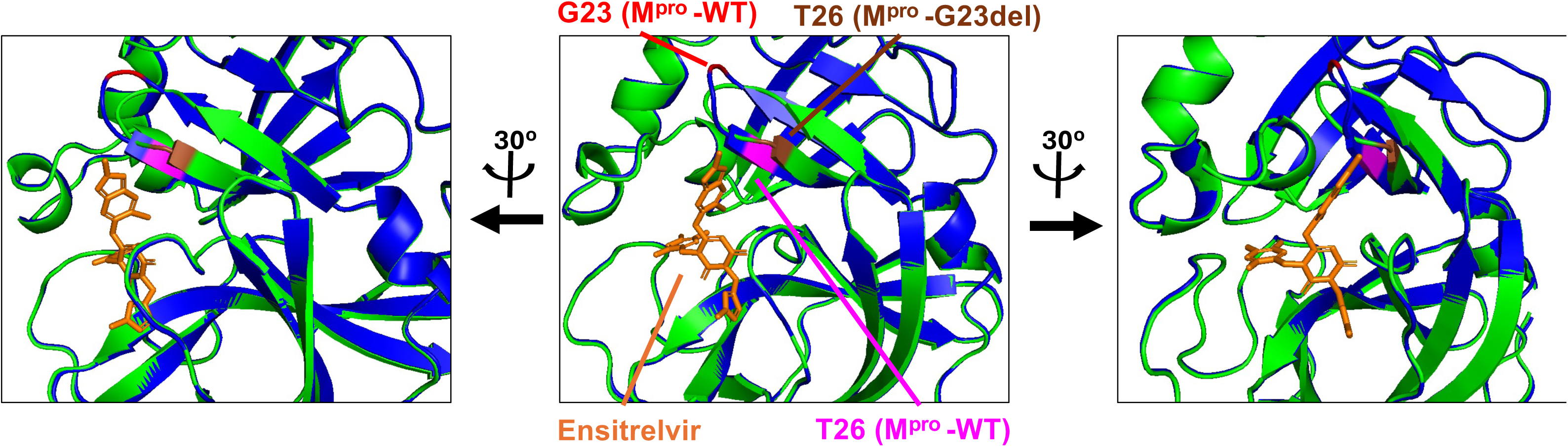
Structural analysis of the G23del in M^pro^. Predicted structures of M□□□-WT (blue) and the M□□□-G23del mutant (green), using AlphaFold2, were aligned and analyzed in PyMOL. Ensitrelvir (orange) forms a hydrogen bond with T26 (magenta) in the M□□□-WT protein. The position of T26 (brown) in the M□□□-G23del mutant is displaced relative to the structure of M□□□-WT. The central panel shows the structural alignment, while the left and right panels present orthogonal views rotated ±30° around the vertical axis to illustrate the conformational changes in the mutant protein.

## DISCUSSION

The capacity of SARS-CoV-2 to evade vaccine-induced immunity, therapeutic antibody biologics, and small-molecule antiviral drugs remains a critical public health challenge (2–5). Neutralizing antibodies administered therapeutically or elicited by mRNA or other vaccines primarily target the spike (S) surface glycoprotein, which has a high mutational degree of freedom (42–45). This plasticity in SARS-CoV-2 S has resulted in the loss of efficacy of previously authorized therapeutic monoclonal antibodies and necessitates continual updates of vaccine formulations (6–13). In contrast, viral targets of antivirals, such as the M^pro^ or RdRp, are evolutionarily more constrained. This conservation forms the foundation for the broad efficacy of such inhibitors against current circulating variants and even across related coronaviruses (14–19). Nevertheless, mutations conferring robust resistance to antiviral drugs have been reported (35, 46–50). Since nirmatrelvir and ensitrelvir target M^pro^, several amino acid substitutions such as E166V and L167F confer cross-resistance to nirmatrelvir and ensitrelvir (33, 51). Importantly, clinical usage of these two antivirals for the treatment of SARS-CoV-2 infections has led to the emergence of M49I, E166A/V, and the combination L50F+E166A mutations (49, 50, 52). Therefore, screening and functional characterization of resistance mutations, together with robust viral surveillance, are imperative for guiding future antiviral strategies.

To address the safety concerns associated with conventional resistance selection analysis using virulent forms of SARS-CoV-2, we developed a luminescent, attenuated recombinant SARS-CoV-2 (Δ3a7b-Nluc WT). Deletion of ORF3a and ORF7b resulted in significant viral attenuation *in vivo* while preserving viral replication. Expression of Nluc enabled sensitive and quantitative tracking of viral replication, including resistance to antiviral treatment (37–39). This platform effectively addressed biosafety concerns associated with resistance profiling of virulent strains, while studying bona-fide replication-competent SARS-CoV-2 to identify and characterize drug resistance mechanisms.

Using this platform, we discovered a novel G23 deletion (G23del) in SARS-CoV-2 M^pro^ that confers resistance to ensitrelvir but not nirmatrelvir. Binding affinity assays confirmed that G23del completely abolished affinity of ensitrelvir for M^pro^, whereas binding to nirmatrelvir was only minimally affected. Structurally, a previous study proved that ensitrelvir docks to M^pro^ through the S1, S2, and S1′ subsites. In the S1’ subsite, ensitrelvir interacts with the NH of main chain T26, which is located in the same β-sheet as G23, via hydrogen bonding of its 6-chloro-2-methyl-2H-indazole moiety. In addition, H163, C145, G143, and Q189 are also engaged in hydrogen-bonds (25). In contrast, nirmatrelvir forms a reversible covalent bond with C145 at the S1′ subsite through its nitrile warhead in P1′, resulting in the formation of a thioimidate adduct that establishes a hydrogen bond with the G143 residue (53, 54). The binding of nirmatrelvir to M^pro^ does not involve the β-sheet in which G23 is located. Structural predictions of the mutated M^pro^ compared to the WT protein posited the primary structural differences between WT and G23del in the β-sheet of the S1′ subsite where G23 is located. This pose is consistent with significantly altered sensitivity of M^pro^-G23del to ensitrelvir, without concomitant change in susceptibility to nirmatrelvir.

In conclusion, utilizing the attenuated SARS-CoV-2 lacking ORFs 3a and 7b proteins and expressing Nluc, we have safely identified and validated the new ensitrelvir-specific resistance mutation G23del. We demonstrated that this mutation confers significant resistance to ensitrelvir *in vitro* and *in vivo* without affecting the ability of nirmatrelvir to inhibit Δ3a7b-Nluc G23del infection. Mechanistic studies revealed that resistance is directly based on reduced binding affinity of ensitrelvir to M^pro^ containing the G23 deletion. Our findings established proof-of-concept for the safe and responsible use of the Δ3a7b-Nluc variant to identify and characterize drug resistant mutants without risk of contributing to the evolution of pathogenic SARS-CoV-2.

## MATERIALS AND METHODS

### Biosafety

All *in vitro* experiments involving the recombinant SARS-CoV-2 Δ3a7b-Nluc WT were performed under Biosafety Level 2+ (BSL-2+) containment laboratories. Experiments to conduct research under Biosafety Level 2 with BSL3 practices (BSL-2+) with SARS- CoV-2 Δ3a7b-Nluc and to identify drug resistant mutants with Δ3a7b-Nluc were approved by the National Institutes of Health Office of Science Policy (NIH OSP). *In vivo* studies were conducted in Animal Biosafety Level 3 (ABSL-3) facilities at the Texas Biomedical Research Institute. These studies were reviewed and approved by the Institute’s Institutional Biosafety Committee (IBC) and Institutional Animal Care and Use Committee (IACUC).

### Cells and Viruses

Vero cells stably expressing human hACE2 and TMPRSS2 (Vero-AT) were obtained from BEI Resources and maintained in Dulbecco’s modified Eagle medium (DMEM) supplemented with 10 µg/ml puromycin (InvivoGen), 10% fetal bovine serum (FBS; VWR), and 100 U/ml penicillin-streptomycin (Corning) at 37°C in a 5% CO_₂_. The Δ3a7b- Nluc WT was previously described (36).

### Isolation of Ensitrelvir Drug-Resistant Mutant (DRM-E)

Vero-AT cells (12-well plates, triplicate wells) were infected with 100 plaque forming units (PFU)/well of Δ3a7b-Nluc WT. After 1 hour of viral adsorption, cells were washed and cultured in medium containing increasing concentration of ensitrelvir (0.05 µM as initial concentration). After 72 hours, cell culture supernatants from wells showing >50% cytopathic effect (CPE) at the highest drug concentration were collected and passaged onto fresh Vero-AT cells using increasing ensitrelvir concentrations. After 10 serial passages, the resulting DRM-E was amplified and stored at -80°C for future use.

### Immunofluorescence Assay (IFA)

Vero-AT cells (6-well plate) were infected with the indicated viruses at a multiplicity of infection (MOI) of 0.001. After 1-hour viral adsorption, cells were incubated with post-infection medium (DMEM supplemented with 2% FBS and penicillin-streptomycin) containing different concentrations of ensitrelvir (0, 0.03, 3, or 300 µM). At 48 hours post-infection, cells were fixed with 10% formalin, permeabilized with 0.5% Triton X-100, and stained using a nucleocapsid (N) protein monoclonal antibody (1C7C7), followed by an FITC-conjugated anti-mouse secondary antibody. Nuclei were counterstained with DAPI. Images were acquired using an EVOS fluorescence microscope (Thermo Fisher Scientific).

### Plaque Reduction Neutralization Test (PRNT)

Confluent Vero-AT cells (96-well plate, quadruplicates) were infected with 100–200 PFU/well of the indicated viruses. After 1 hour of viral adsorption, the inoculum was replaced with post-infection medium containing serial dilutions of ensitrelvir (starting concentration of 400 µM), nirmatrelvir (starting concentration of 100 µM) or remdesivir (starting concentration of 100 µM) and 1% Avicel. At 18 hours post-infection, cells were fixed with 10% formalin, permeabilized with 0.5% Triton X-100, and immunostained as described above using the 1C7C7 monoclonal antibody. Detection was performed using the Vectastain ABC-HRP kit and DAB Substrate Kit (Vector Laboratories). Plates were imaged by Bioreader 7000 F-z (BIOSYS), and plaque counts were used to calculate the half-maximal effective concentration (EC_₅₀_).

### Sequencing

Viral genome sequences were confirmed by whole-genome sequencing using the MinION platform (Oxford Nanopore Technologies). Total RNA was extracted from infected Vero-AT cells using TRIzol reagent (Thermo Fisher Scientific) according to the manufacturer’s instructions. The cDNA was synthesized from extracted RNA using the SuperScript IV VILO Master Mix (Invitrogen). Subsequent PCR amplification was performed using the ARTIC Network nCoV-2019 version 5.3.2 primer panel (Integrated DNA Technologies). Sequencing libraries were prepared with the Native Barcoding Kit 24 (v14; SQK-NBD114.24, Oxford Nanopore Technologies) following the manufacturer’s protocol. Sequencing was carried out on R10.4.1 flow cells (FLO-MIN114, Oxford Nanopore Technologies). Raw reads were base-called and mapped to the reference genome using Geneious Prime software for consensus sequence generation and variant analysis. To confirm the presence of the G23del mutation, the region encoding NSP5 was subsequently amplified by PCR using the Expand High Fidelity PCR System (Sigma-Aldrich) from cDNA samples. The resulting amplicons were purified and subjected to Sanger sequencing by Plasmidsaurus Inc.

### Recombinant viruses

The recombinant SARS-CoV-2 Δ3a7b-Nluc WT containing the G23del mutation was rescued in Vero-AT cells using a previously established protocol (55, 56). Briefly, a bacterial artificial chromosome (BAC) harboring the full-length viral genome of Δ3a7b-Nluc WT containing the G23del was transfected into confluent monolayers of Vero-AT cells (6-well plate) using Lipofectamine 2000 (Thermo Fisher Scientific). At 24 hours post-transfection, the medium was replaced with post-infection medium. At 48 hours post-transfection, Vero-AT cells were subsequently harvested and transferred to T75 flasks and incubated with post-infection media for an additional 72 hours. Finally, cell culture supernatants containing the rescued virus were harvested and stored at -80°C.

### Plaque Assay and Immunostaining

Confluent monolayers of Vero-AT cells (6-well plate) were infected with 10-fold serial dilutions of the indicated viruses for 1 hour at 37°C. Following adsorption, the virus inoculum was removed, and cells were overlaid with DMEM containing 1% agarose. After 96 hours of incubation at 37°C, cells were fixed with 10% formalin. For luminescence-based plaque visualization, agar overlays were removed, and cells were incubated with the Nano-Glo Luciferase Assay Substrate (Promega) according to the manufacturer’s instructions. Plates were imaged using a ChemiDoc MP Imaging System (Bio-Rad). For immunostaining, cells were permeabilized with 0.5% Triton X-100 followed by incubation with the 1C7C7 N protein monoclonal antibody. Detection was performed using the Vectastain ABC-HRP kit and DAB Substrate Kit (Vector Laboratories) by ChemiDoc MP Imaging System (Bio-Rad). Cells were counterstained with crystal violet and imaged by ChemiDoc MP Imaging System (Bio-Rad).

### Viral Growth Kinetics and Nluc assay

Confluent monolayers of Vero-AT cells (6-well plate, triplicates) were infected at a multiplicity of infection (MOI) of 0.01. After 1 hour of viral adsorption at 37°C, cells were washed with PBS and maintained in post-infection medium. Cell culture supernatants were collected at 0-, 24-, 48-, and 72-hours post-infection. Viral titers were determined by plaque assay and immunostaining as described above. Secreted Nluc activity was quantified using the Nano-Glo Luciferase Assay System (Promega).

### Animal experiments

Five-week-old female K18-hACE2 transgenic mice were obtained from The Jackson Laboratory and housed under specific pathogen-free conditions at the Texas Biomedical Research Institute animal facility for 7 days before infection. Mice were anesthetized via intraperitoneal injection of ketamine and inoculated intranasally with 10^7^ PFU of either Δ3a7b-Nluc WT or the Δ3a7b-Nluc G23del mutant virus (n = 3 per group) at day 0. At 1- day post-infection, mice were assigned to treatment or control groups. The treatment group (n = 3) received ensitrelvir resuspended in 0.5% methylcellulose, at a dose of 60 mg/kg via oral gavage twice daily until 3-day post-infection. Control mice (n = 3) received vehicles alone (0.5% methylcellulose) on the same schedule. At 4 -day post-infection, mice were anesthetized with ketamine and retro-orbitally injected with 100 µL of Nano-Glo Luciferase Substrate diluted 1:10 in PBS. Then, animals were euthanized by intraperitoneal injection of Fatal Plus. Lungs were immediately harvested and analyzed using an Ami HT *in vivo* imaging system (Spectral Instruments) to quantify Nluc-derived luminescence. Following imaging, both lungs and nasal turbinate were collected for subsequent viral titration via plaque assay on Vero-AT cells.

### Expression, and purification of SARS-CoV-2 M^pro^-WT and M^pro^-G23del mutant

The gene encoding SARS-CoV-2 M^pro^-WT and M^Pro^-G23del were cloned into the pET28b-smt3 plasmid. Both WT and G23del M^pro^ were expressed and purified using the same protocol. Briefly, pET28b-smt3 plasmids were transformed into *E. coli* BL21 Nico cells (New England Biolabs) and large cultures (Terrific Broth medium) were grown at 37°C. Protein expression in these cultures was induced by adding 0.4 mM Isopropyl β- D-1-thiogalactopyranoside at an O.D. of 0.6 - 0.8 and growing at 18 °C overnight. The bacteria were pelleted down by centrifugation at 6,000 rpm for 20 minutes. The pellets were resuspended in lysis buffer (25 mM Tris, pH 8.0, 0.5 M NaCl, 10% Glycerol, 0.2 mM TCEP, 5 mM Imidazole, supplemented with DNase I, PMSF, and one tablet of protease inhibitor cocktail (Roche)) and lysed using a microfluidizer. Bacteria lysates were centrifuged at 40,000 rpm for 45 minutes and filtered through a 0.22 µm filter. Supernatants were loaded onto a 5 mL Ni-NTA affinity column (EconoFit Nuvia, BioRad). Colums were washed with washing buffer (25 mM Tris, pH 8.0, 0.5 M NaCl, 10% Glycerol, 0.2 mM TCEP, 5 mM Imidazole) and proteins were eluted using an elution buffer (25 mM Tris, pH 8.0, 0.5 M NaCl, 10% Glycerol, 0.2 mM TCEP, 1 M Imidazole). Eluted proteins were treated with recombinant Ulp-1 protease to cleave off the His_6_-SUMO tag, and the tag was separated by a subsequent passage through a Ni-NTA affinity column. Proteins were further purified by HiTrap Q HP column (Cytiva) using a buffer (25 mM Tris, pH 8.0, 10% Glycerol, 0.2 mM TCEP) with a linear gradient of salt from 40 mM to 2.0 M. Proteins were finally purified by a HiLoad 16/600 Superdex 75 pg column (Cytiva) in a buffer containing 20 mM Tris, pH 7.8, 0.15 M NaCl, 1 mM EDTA, and 1 mM DTT. Purified proteins (>95% purity) were concentrated to 20 mg/mL and used immediately or flash-frozen in liquid nitrogen, and stored at -80°C.

### Quantitative binding affinity

The isothermal titration calorimetry (ITC) experiments were performed using the MicroCal PEAQ-ITC instrument (Malvern Panalytical). The proteins were extensively dialyzed in a buffer containing 20 mM Tris, pH 7.8, 0.1 M NaCl, and 1 mM TCEP prior to ITC measurements. After cleaning the sample cell and injection needle, 20 or 30 µM of M^pro^-WT and M^pro^-G23del proteins were loaded into the sample cell using a micro- syringe. Then, 0.2 or 0.3 mM ensitrelvir or nirmatrelvir antivirals were loaded into a 40 μL titration syringe. A total of 1.5 µL of each ligand was injected 25 times with 150 seconds of spacing between injections. The experiments were performed at 25°C. The deionized water was injected into the reference cell as a heat-balance control. After 25 titrations of ensitrelvir or nirmatrelvir, the data were fitted using a single-site binding model to calculate stoichiometry (N) and affinity (K_d_) binding.

### Structure Prediction

The amino acid sequences of M^pro^-WT and M^pro^-G23del were submitted to AlphaFold2 to generate structural predictions (57). The resulting models were structurally aligned and analyzed using PyMOL (The PyMOL Molecular Graphics System, Version 2.5 Schrödinger, LLC) to analyze the structural perturbations induced by the deletion.

### Statistical analysis

All data are presented as the mean ± standard deviation (SD). Statistical analyses were performed using GraphPad Prism software. For comparisons across multiple groups, a one-way analysis of variance (ANOVA) was conducted, followed by Tukey’s post hoc test for multiple comparisons. A p-value of less than 0.05 was considered statistically significant.

## ACKNOWLEDGMENTS

We thank BEI Resources for providing Vero-AT cells. This work was supported, in part, by public health service grant U19 AI171403 (to R.K.P.), R01 AI161363, and R01 AI161363-03S1 (to Y.K.G. and L.M.S.) from the NIH/NIAID. This work was also partly supported by CRIPT (Center for Research on Influenza Pathogenesis and Transmission), a NIAID-funded Center of Excellence for Influenza Research and Response (CEIRR, contract #75N93021C00014) to A.G.-S.

---

**Supplementary Figure S1. Binding isotherms of Ensitrelvir and Nirmatrelvir.** ITC experiments with M^pro^-WT and M^pro^-G23del were performed using 30 µM protein and 0.3 mM of ligand at 25°C in identical experimental conditions. The upper panel illustrates the raw heat signals while the lower panel shows the integrated heat and the fit using a one-site binding model for Ensitrelvir with M^pro^-WT (A), Ensitrelvir with M^pro^-G23del (B), Nirmatrelvir with M^pro^-WT (C), and Nirmatrelvir with M^pro^-G23del (D).

