## Supplementary figures and images for "Identification and characterization of a SARS-CoV-2 M^pro^ G23 deletion ensitrelvir-resistant mutant"

### Supplemental Figure 1

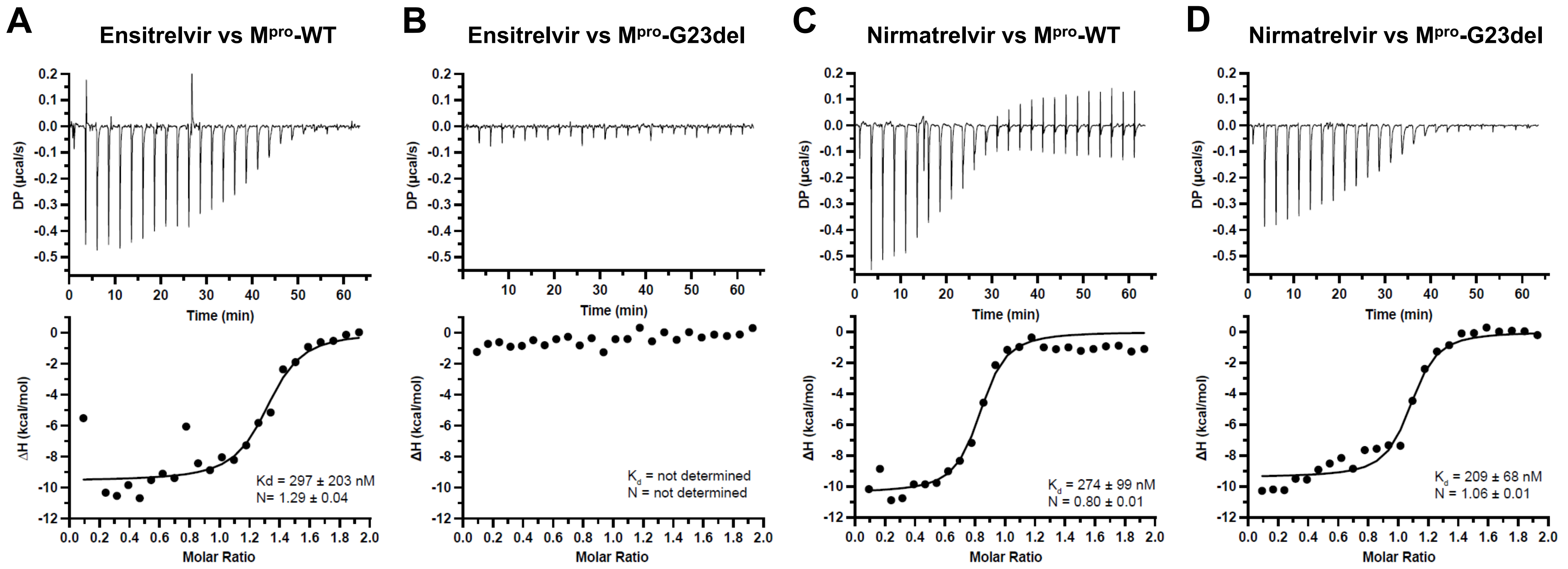
